# Regions of Highly Recurrent Electrogram Morphology at Sites of Low Cycle Length Accurately Reflect Arrhythmogenic Substrate for Atrial Fibrillation – Implications For a New, Mechanism Guided Therapeutic Approach for Atrial Fibrillation

**DOI:** 10.1101/2020.12.10.419754

**Authors:** Shin Yoo, Markus Rottmann, Jason Ng, David Johnson, Bassel Shanab, Anna Pfenniger, Gail Elizabeth Geist, Suman Mandawa, Amy Burrell, Wenwei Zhang, J Andy Wasserstrom, Bradley P Knight, Rod Passman, Jeffrey J Goldberger, Rishi Arora

## Abstract

**Background:** Although atrial electrograms (EGMs) are thought to reflect pathophysiological substrate for atrial fibrillation (AF), it is not known which electrograms are suitable targets during AF ablation. We hypothesized that electrogram morphology recurrence (EMR) better reflects arrhythmogenic AF substrate than traditional frequency and complexity measures of AF. In a canine rapid atrial pacing (RAP) model of AF, we assessed the relationship between EMR and traditional AF electrogram measures, rotational activity in the atria, fibrosis, myofiber orientation and parasympathetic innervation.

**Methods:** Persistent AF was induced in 13 dogs by RAP for 6-8 weeks. High-density epicardial mapping (117 electrodes) was performed in six atrial sub-regions. EMR measures Recurrence percentage (Rec%) and cycle length of the most frequent electrogram morphology (CL_R_), Fractionated Interval (FI), Organization Index (OI), Dominant Frequency (DF) and Shannon’s Entropy (ShEn) were analyzed before and after atropine administration. Myocyte fiber orientation, amount of fibrosis and spatial distribution of parasympathetic nerve fibers were quantified.

**Results:** Rec% was greatest in the appendages, and CL_R_ was lowest in the posterior left atrium. Rec%/CL_R_ correlated with FI, OI and the complexity measure ShEn, but not with DF. All electrogram measures were poorly correlated with fibrosis and myofiber anisotropy. Rec% correlated closely with stability of rotational activity. Unlike other measures, Rec% correlated closely with spatial heterogeneity of parasympathetic nerve fibers; this was reflected in CL_R_ response to atropine.

**Conclusion:** EMR correlates closely with stability of rotational activity and with the pattern of atrial parasympathetic innervation. CL_R_ may therefore be a viable therapeutic target in persistent AF.

## Introduction

AF drivers such as rotational and focal activities are thought to sustain AF and are often located near pulmonary veins (PVs).^1, 2^ PV isolation has been widely applied to treat patients with AF, however, the suboptimal ablation outcomes in patients with persistent AF (< 50%) suggests that other atrial regions besides the PVs may be responsible for sustaining AF activity.^3, 4 5^ Multiple electrogram based approaches have been promulgated to help identify regions of interest, including mapping complex fractionated atrial electrograms (CFAE), dominant frequency (DF), Shannon’s entropy (ShEn), voltage, and focal impulse and rotor mapping (FIRM). None have been established as a widely successful strategy. The challenges of using DF analysis for AF electrograms have been shown^6, 7^, underscoring the need for a new approach.

There are multiple factors that determine atrial electrograms in AF, including characteristics of the recording electrodes, the activation rate, the underlying electrophysiologic properties, and the underlying tissue anatomy/pathology. Analysis of the electrogram morphology may provide information on the latter 3 components. Indeed, electrogram morphology recurrence analysis can provide important classification of electrograms.^8^ We applied electrogram morphology recurrence analysis in a preliminary clinical study of patients with AF^9^. In that study, multisite mapping of the right and left atrium was performed. At each site, the recurrence percentage of the most frequent electrogram morphology was determined (Rec%), as well as the cycle length of this most frequent electrogram morphology (CL_R_). None of the patients with shortest CL_R_ in the right atrium had a successful outcome from left atrial ablation. Rec% provides a measure of the consistency of electrogram morphology. This is expected to be high near AF drivers due to the consistency of activation, but may also be high at other sites for anatomic reasons. Hence, the CL_R_ provides an assessment of the sites with both consistent and rapid activation.

We hypothesize that measures of electrogram morphology recurrence – Rec% and/or CL_R_ – are a more sensitive marker of pathophysiological substrate for AF than traditional electrogram measures of AF. Indeed, even though several investigators have suspected that potential AF mechanisms such as heterogeneous myocyte fiber orientation, ion channel remodeling, structural remodeling (i.e. fibrosis)^10–12^ and altered parasympathetic nervous system signaling^13^ may contribute to AF electrogram formation,^14^ very few studies have systematically examined the relationship between AF electrogram measures and the underlying structural and molecular aspects of AF substrate. The present study was therefore designed to obtain a thorough assessment of the electrophysiological and structural basis of AF electrograms – both established frequency and complexity parameters and the recurrence morphology measures - by performing detailed, high resolution epicardial mapping in all sub-regions of the left and right atrium in a canine model of rapid atrial pacing (RAP) induced persistent AF. The specific goals of this study were as follows: a) to assess the regional and sub-regional distribution of established measures of AF frequency and complexity (DF, OI, FI and ShEn) and our novel recurrence morphology analyses; b) to determine the precise relationship between these different measures; c) to determine whether there is a relationship between electrophysiological substrate for AF – specifically the ability of the atria to harbor rotational (reentrant) activities – and recurrence morphology; d) to assess whether myocyte fiber orientation and fibrosis affect recurrence morphology and e) to determine the relationship between parasympathetic nerve innervation and recurrence morphology.

## Methods

### Rapid atrial pacing model

Thirteen purpose-bred hounds weighing 25 to 35 kg over one year in age were used for this study. The dogs underwent rapid atrial pacing (RAP) for the AF electrogram mapping. The RAP model for AF was performed similarly to previously published techniques.^15^ Sterile surgery for pacemaker implantation was performed for each dog. Endocardial pacing leads were placed into the right atrial appendage (RAA). The pacemakers were then programmed to pace at 600 bpm at a minimum of four times the capture threshold. The dogs were paced for 6 to 8 weeks to induce sustained AF. Once persistent/sustained AF had been induced, the animals were subjected to a terminal, open-chest electrophysiological mapping study. The animal study protocol conforms to the Guide for the Care and Use of Laboratory Animals published by the U.S. National Institutes of Health (NIH Publication No. 85-23, revised 1996) and was approved by the Animal Care and Use Committee of Northwestern University. Before undergoing pacemaker implantation and electrophysiological mapping, all animals were premedicated with acepromazine (0.01 – 0.02 mg/kg) and were induced with propofol (3-7 mg/kg). All experiments were performed under general anesthesia (inhaled) with isoflurane (1-3 %). Adequacy of anesthesia was assessed by toe pinch and palpebral reflex.

### *In vivo* electrophysiological mapping

#### AF Electrogram Mapping

At the terminal study, high-density epicardial activation mapping was performed using the epicardial UNEMAP mapping system (Univ. of Auckland, Auckland, New Zealand) containing 130 electrodes (inter-electrode distance of 2.5 mm) and 117 bipolar EGM recordings at 1 kHz sampling rate on a triangular plaque. Consecutive recordings were obtained using the GE Prucka Cardiolab system (GE Healthcare, Waukesha, WI, USA). We collected bipolar electrograms from six different regions and in quadrants of these regions in the left atrial free wall (LAFW), posterior left atrium (PLA), left atrial appendage (LAA), posterior right atrium (PRA), right atrial free wall (RAFW), right atrial appendage (RAA). AF EGMs were recorded for calculating the following EGM parameters: 1) Recurrence Percentage (Rec%), 2) Recurrence Cycle Length (CL_R_), 3) Dominant Frequency (DF), 4) Organization Index (OI), 5) Fractionation Interval (FI) and 6) Shannon’s Entropy (ShEn).

#### Recurrence Percentage (Rec%)

EMR analysis of AF involves the creation of morphology recurrence plots, as previously described^16^ with the modification of a paper by Eckmann et al.^17^ Rec% is defined as the percentage of the most common morphology and was recently developed by our group for AF electrogram analysis.^18^ The EGM morphology recurrence plots of AF recordings at each site were calculated by cross-correlation of each detected activation using an iterative method^18^ with all the other activations during a 10 s recording period (examples in Fig 1A). Rec% was then computed as percentage of the number of the most common morphology against the total number of activations.^16^ Electrograms with a high degree of similarity have cross-correlation values near 1, while dissimilar electrograms had values closer to zero.

**Figure 1.**
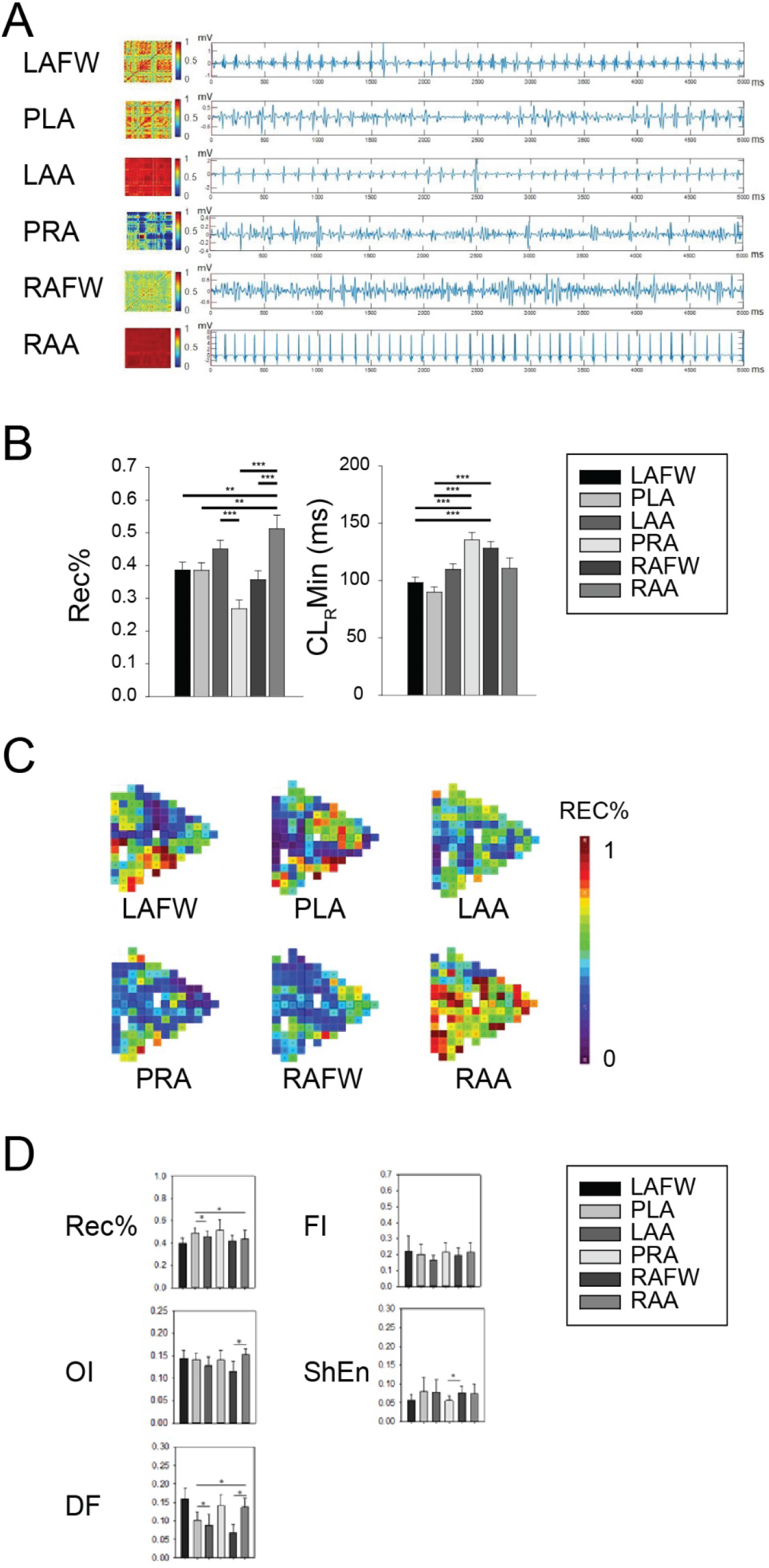
A, Illustration of a color coded cross-correlation matrix of all activations (left) and EGM signals (right) in six atrial regions. B, Regional distribution of Rec% and CL_R_Min in six atrial regions. Data are presented as mean ± SEM; ** p < 0.01 and *** p < 0.001. C, Examples of spatial distribution of Rec% in six atrial regions. D, Coefficient of variation of Rec%, FI, OI, ShEn and DF in six atrial regions. Data are presented as mean ± SEM; * p < 0.05. one-way ANOVA with Holm-Sidak method for pairwise multiple comparison.

#### Cycle length of the most recurrent morphology (CL_R_)

Cycle length of the most recurrent morphology was obtained by dividing the average cycle length for all EGMs by Rec%.

#### Dominant Frequency (DF)

DF is a frequency domain measure of the activation rate. The EGM signals recorded during a 10 s period were bandpass filtered with cutoff frequencies of 40 and 250 Hz and rectification. The power spectrum of the EGM segment was computed using the Fast Fourier transform. The frequency with the highest power in the power spectrum was considered the DF.

#### Organization Index (OI)

OI is a frequency domain measure of temporal organization or regularity^19 20^. OI was calculated as the area under 1-Hz windows of the DF peak and the next three harmonic peaks divided by the total area of the spectrum from 3 Hz up to the fifth harmonic peak. It has been shown that AF episodes with recordings with high OI are more easily terminated with burst pacing and defibrillation.^20^

#### Fractionation Interval (FI)

FI is the mean interval between deflections detected in the EGM segment.^21^ Deflections were detected if they meet the following conditions: 1) the peak-to-peak amplitude was greater than a user-determined noise level, 2) the positive peak was within 10 ms of the negative peak, and 3) the deflection was not within 50 ms of another deflection. The noise level was determined by selecting the amplitude level that would avoid the detection of noise-related deflections in the iso-electric portions of the signal. FI is dependent on both the AF cycle length and the fractionation of the EGM.

#### Shannon’s Entropy (ShEn)

ShEn is a statistical measure of complexity.^22^ The 4000 or 3908 (depending on the 1kHz or 977 Hz sample rate) amplitude values of each EGM segment were binned into one of 29 bins with a width of 0.125 standard deviations. ShEn was then calculated as:

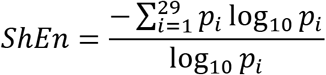

In this equation, pi is the probability of an amplitude value occurring in bin *i*.

#### Parasympathetic Blockade

Parasympathetic blockade was performed by atropine (0.04 mg/kg; Med-Pharmex Inc). AF characteristics were analyzed at baseline, and after parasympathetic blockade.

#### Detection of Reentries and their Stability

Reentries were detecting based on phase singularity detections using the Hilbert phase and sinusoidal recomposition.^23^ The phase φ(t) of the complex-valued analytic signal z(t) was calculated as with the real and imaginary part of the analytic sigal:

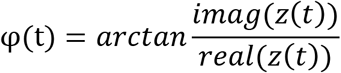

The stability was assessed as duration of observed reentrant activity over time.

### Tissue preparation for analysis

After mapping the epicardial surface, we excised the heart out of the chest and immersed in ice-cold cardioplegia solution containing (in mmol/l) NaCl 128, KCl 15, HEPES 10, MgSO_4_ 1.2, NaH_2_PO_4_ 0.6, CaCl_2_ 1.0, glucose 10, and heparin (0.0001 U/ml); pH 7.4. All solutions were equilibrated with 100% O_2_. We cannulated the heart via the aorta and perfused with ice-cold cardioplegia solution containing protease inhibitors (Millipore Sigma, P8340) until the vessels were clear of blood, and tissue was cold. Atrial tissue was excised and 6 atrial regions were dissected. Specimens from 4 dogs were fixed in 10% formalin and embedded in paraffin for further examination.

### Masson’s trichrome staining

Paraffin sections with 5 μm thickness were stained using Masson’s Trichrome stain kit (Sigma) as described previously.^24^ In brief, paraffin was removed in Xylene and the sections were then rehydrated with ethanol series. The sections were treated with Bouin’s mordant at room temperature overnight. The following day the sections were stained in Weigert’s Iron Hematoxylin solution and Beibrich Scarlet-Acid fuchsin. Then the sections were incubated in the phosphomolybdic-phosphotungstic acid solution. The sections were stained in the Aniline Blue solution. The sections were incubated in 1% Glacial acetic acid, were dehydrated through ethanol series and were placed in Xylene, A coverslip was placed using cytoseal mounting media on the sections.

### Examination of uniformity of fiber orientation and degree of fibrosis

Masson’s trichrome stained tissue sections at different depths (0, 200, and 500 μm) from the epicardial surface were digitized with the NanoZoomer 2.0-HT at 5x magnification. For quantitative morphometric analysis, the whole sections were divided into quadrant by drawing regions of interest. In order to quantify uniformity of fiber orientation in RAP atrial tissue sections at 200 μm and 500 μm from the epicardial surface, we were able to automatically quantify the orientation of the myocytes using an ImageJ plugin called FibrilTool.^25^ Fibril-Tool calculates a value referred to anisotropy index (AI), which is a measure of how parallel the fibers are with respect to each other. We further analyzed the degree of fibrosis in Masson’s trichrome stained tissue section at 200 μm levels using ImageJ with a macro^26^ as there was qualitative similarity among sections at different depths.

### Quantification of parasympathetic nerve fiber density by immunohistochemistry

Cryosections from 6 atrial regions (LAFW, PLA, LAA, PRA, RAFW, and RAA) taken from −80 °C freezer were air-dried and underwent fixation with 75% acetone/25% ethanol and washed in Tris-buffered saline with 0.5% tween 20 (TBS-T). The sections were then treated in 3% hydrogen peroxide. After washing in TBS-T, the sections were blocked in protein block reagent (Dako) and then incubated overnight with anti-mouse acetylcholinesterase (AChE, Millipore-Sigma, MAB303) at 4 °C. The next day, the sections were washed in TBS-T and incubated with HRP-conjugated anti-mouse secondary antibody (Dako, K4000). The sections were stained brown by incubation of 3,3′-diaminobenzidine (DAB). After reapplication of protein block, anti-rabbit antibody for dopamine β-hydroxylase (DBH; Chemicon, AB1538) was incubated for 1 hour at room temperature. After being washed in TBS-T, the sections were incubated with HRP-conjugated anti-rabbit secondary antibody (Dako, K4002) for 30 minutes at room temperature. The sections were stained blue by incubation of 5-bromo-4-chloro-3-indolyl phosphate (BCIP). Cell nuclei were counter-stained in methyl green (Dako). Specimens were then dehydrated in series of ethanol and xylene, mounted in cytoseal mounting media (Thermo Fisher Scientific). Stained sections were examined using transmitted light microscope (Olympus) or TissueFax system (TissueGnostics). Parasympathetic fiber density in the myocardium in acquired whole scan images were quantified by TissueFax system and histoquest software (TissueGnostics).

### Statistical analysis

All values were expressed as mean ± standard error if samples were normally distributed. If normality test was not passed, Box and Whiskers plot was employed. When comparing EGM parameters or tissue characteristics between atrial regions, one-way ANOVA was performed with Holm-Sidak method for pairwise multiple comparison if samples were normally distributed.

If normality test was not passed, Kruskal-Wallis one way ANOVA with Turkey test for pairwise multiple comparison was performed. Effect of atropine on EGM parameters was determined by paired t-test with a Wilcoxon rank sum test. Correlation of two variables was performed with the Pearson Product Moment Correlation. The p values were considered significantly different at p < 0.05.

## Results

In this study, we induced persistent AF in 13 dogs. We analyzed the novel EMR measures - Rec% and CL_R_ - in six different regions in the left and right atrium and compared them with established AF source EGM measures i.e. FI, OI, DF and ShEn. Next, we assessed whether Rec% and CL_R_ can help predict the presence and stability of rotational (reentrant) activity in the fibrillating atrium. We further investigated the effect of degree of fibrosis and myofiber orientation on morphology recurrence. Since parasympathetic nerve remodeling^27–31^ has been shown to be an important mechanism underlying AF, we also investigated the effect of parasympathetic innervation – and parasympathetic blockade – on EMR.

### EMR analysis in canine RAP of model of AF

We analyzed the electrogram morphology in the different atrial regions during AF. Supplemental Figure 1A demonstrates examples of cross-correlation of detected activation waveforms to generate EMR plots. The N x N cross-correlation values are plotted in a two-dimensional color-coded map as shown in Supplemental Figure 1B, where N is the number of activations. In this plot, the x-axis and y-axis represent the first and the second activation template that is cross-correlated. The points in red represent the combination with the highest cross-correlation values near 1, while the points in blue represent the cross-correlation values near 0. The checkerboard pattern shown in Supplemental Figure 1B suggests there is a dominant morphology that periodically recurs for the duration of the recording.

### CL_R_ is lower in the PLA than in other regions of the left and right atrium

We quantified Rec% in the different atrial regions of the canine RAP model of AF. We also calculated the mean cycle length of the most recurrent electrogram morphology (i.e. CL_R_) because there may be areas with very recurrent electrogram morphology that may be too slow to be likely drivers for AF. For this reason, in addition to quantifying the Rec% at a particular site, CL_R_ also needs to be determined.

Figure 1A shows examples of recurrence plots and electrograms in each sub-region. Distinct checkerboard patterns in the different sub-regions indicate that the activation patterns have different levels of complexity. High-density mapping data revealed that Rec% and CL_R_ are regionally variable (Figure 1B). The highest overall Rec% was measured in the appendages and the lowest overall Rec% was observed in the PRA. CL_R_ was lower in the left atrium as compared to the right atrium, with the lowest CL_R_ measured in the PLA.

Since spatial heterogeneity of electrophysiological characteristics is an important predictor of arrhythmogenic substrate, we also analyzed the spatial distribution of Rec% in all mapped regions. The coefficient of variation of Rec% showed significant inter-regional variability, being more homogeneous in the LAA as compared to other atrial regions (Figure 1C and 1D).

### Rec% and CL_R_ are more closely correlated with Fractional Interval and Shannon’s Entropy than with Dominant Frequency

Next, we examined the precise correlation between Rec% and each of the rest of the electrogram measures at each electrode within each mapped region. The methodology for this correlation analysis is shown in figure 2A. As shown in figure 2B, R was ≥ 0.5 in nearly all regions for Rec%-FI and Rec%-ShEn, with Rec%-OI being close to 0.5 in several regions. In distinct contrast Rec%-DF was markedly lower than 0.5 in every region, indicating a very poor correlation between Rec% and DF. Although correlations were somewhat weaker for CL_R_, the correlation between CL_R_ – FI and CL_R_ – ShEn was ≥ 0.5 in most regions. Taken together, Rec% and CL_R_ are moderately correlated with fractionation and complexity measures (FI and ShEn) of AF.

**Figure 2.**
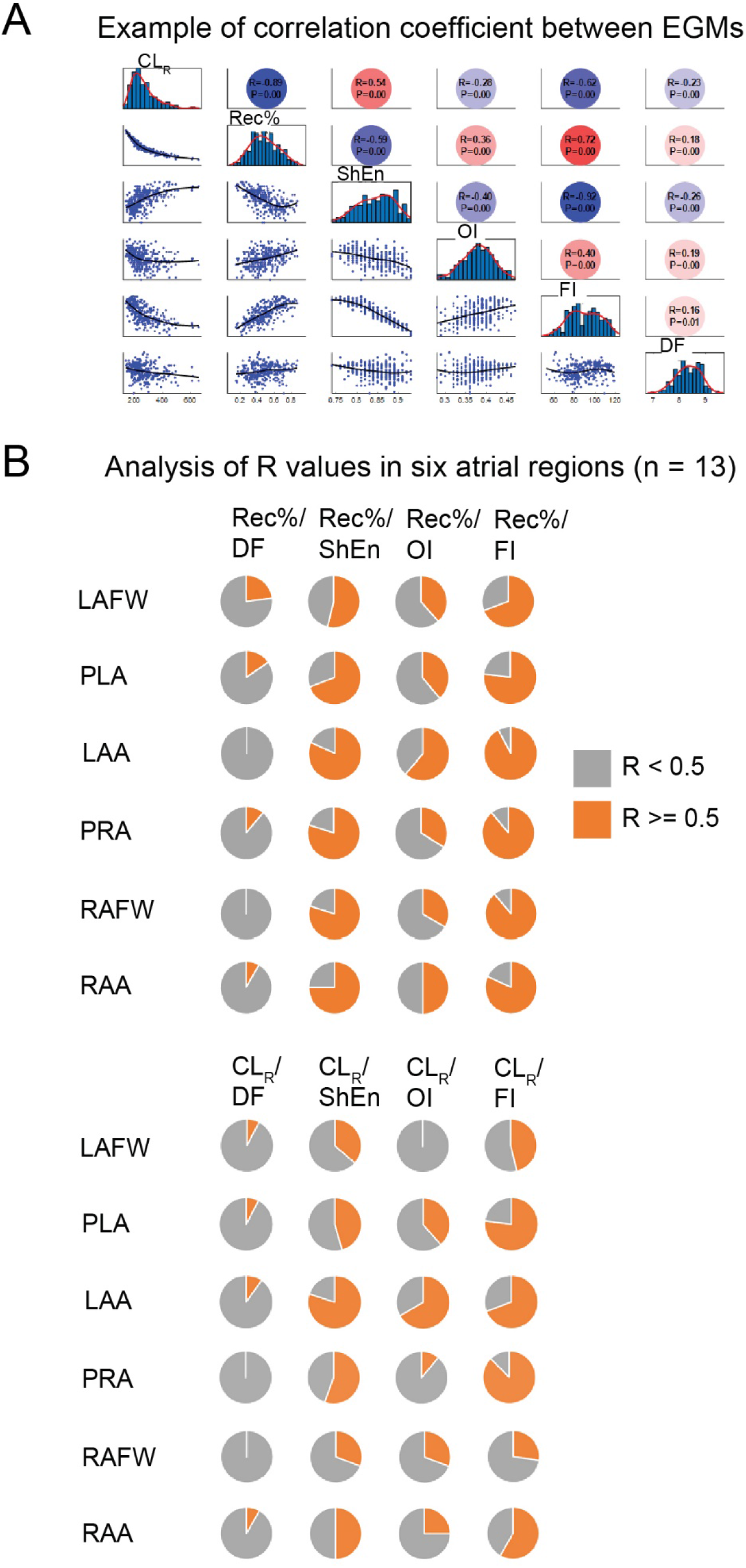
A, Example of calculation of correlation coefficient between EGMs in animal 1 in LAA. B, Analysis of R values of Rec% (top) and CL_R_ (bottom) with other EGMs in six atrial regions. Pie charts show proportion of animals with correlation factor R >= 0.5 (orange) and R < 0.5 (gray).

### Rec% strongly reflects stability of rotational activities in the fibrillating atrium

Evidence of the presence of rotational activities and the characteristics of these rotational activities have previously been shown to correlate with arrhythmogenic substrate in patients with AF. We therefore examined AF sources showing 360 degree rotations in the local activation time maps in all 6 regions (figure 3). Even though reentry was seen in all regions, the temporal stability of confirmed 360 degree rotations (reentries) differed significantly amongst the regions (figure 3A and 3B). The stability of these reentries corresponded closely to that of the regional distribution of Rec%, with the rotations being most stable in the RAA, followed by the LAA, and then the rest of the left and right atrium (figure 3B). The stability of rotational activities correlated closely with Rec% (figure 3C).

**Figure 3.**
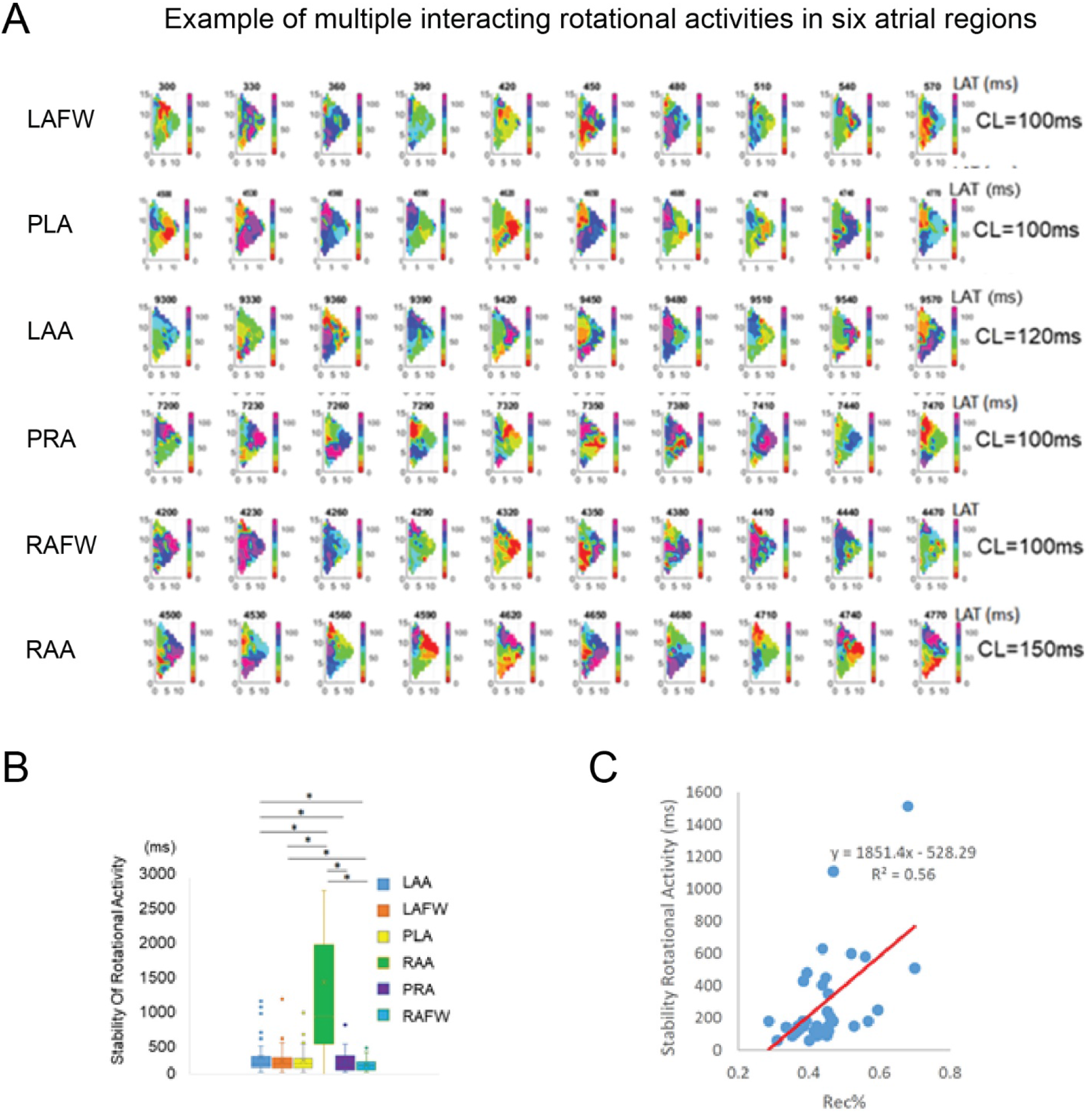
A, Examples of multiple interacting rotational activities in six atrial regions. B, Comparison of stability of rotational activities in six atrial regions. Data are presented in Box and Whiskers plot; * p < 0.05. one-way ANOVA with Holm-Sidak method for pairwise multiple comparison. C, correlation of stability of rotational activities with Rec%.

Since Rec% appears to a highly sensitive marker of stability of reentry, we postulate that the cycle length of the most recurrent morphology (CL_R_) may be highly indicative sites of potential ‘driver’ rotors (see Discussion).

### Rec% and CL_R_ are poorly correlated with fibrosis

A major tissue characteristic thought to affect the number of potential wavefront directions is fibrosis.^32^ Therefore, we quantified the extent of fibrosis in each region and assessed the relationship between fibrosis and AF electrogram characteristics. Figure 4A shows representative images from Masson’s Trichrome stained sections. The amount of fibrosis was regionally variable, being highest in RAFW and lowest in the two appendages. (Figure 4B). There was a modest inverse correlation between fibrosis and FI and between fibrosis and OI (Figure 4C). There was a non-significant trend towards a correlation between fibrosis and Rec% (and between fibrosis and CL_R_).

**Figure 4.**
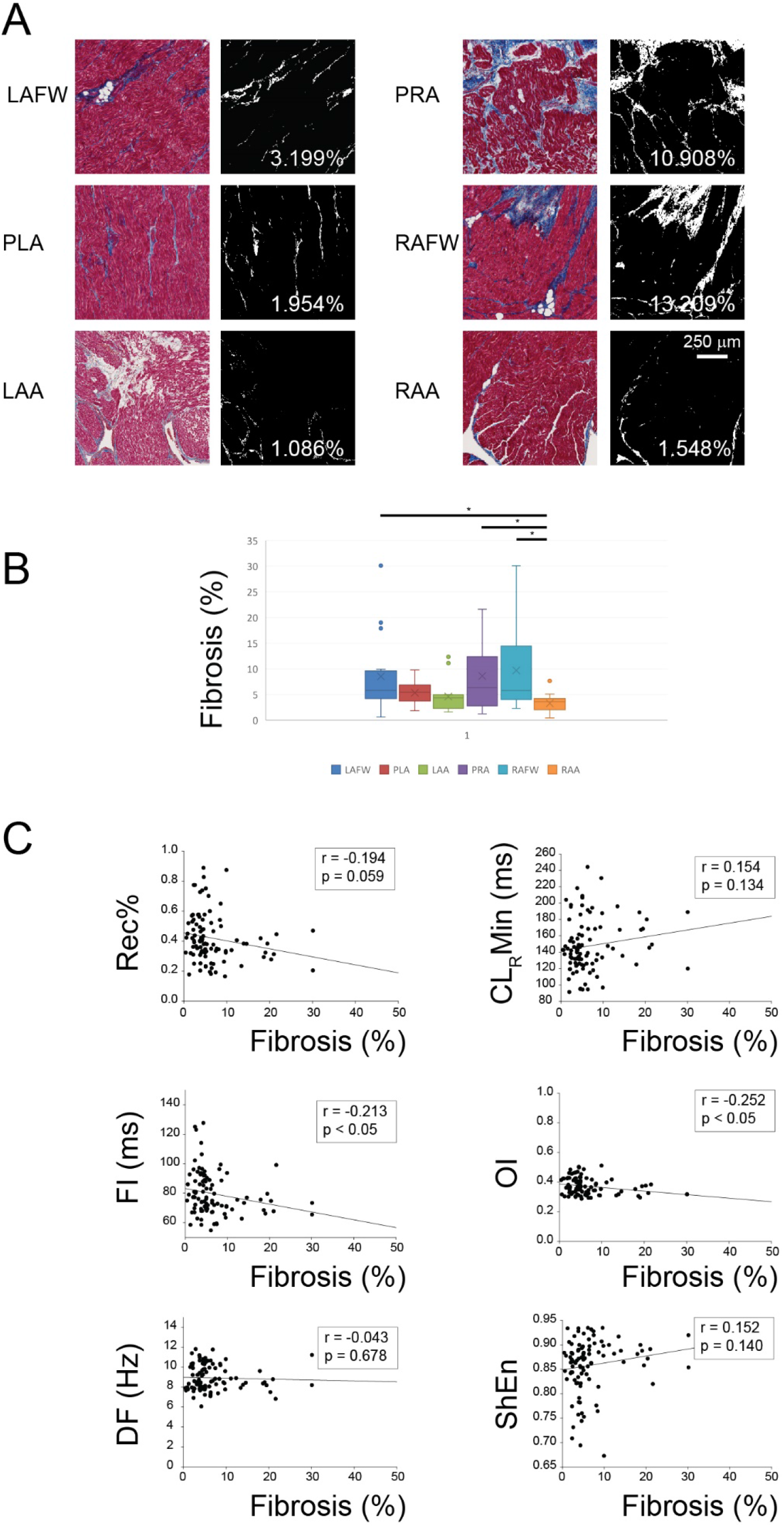
A, Exemplary images of masson’s trichrome stained tissue section and outcome of analysis in six atrial regions. Red indicated myocardium and blue indicated fibrosis. B, Regional differences in fibrosis. Data are presented as mean ± SEM; * p < 0.05. Kruskal-Wallis One Way Analysis of Variance on Ranks with Turkey test for all pairwise comparison. C, Correlation of fibrosis with Rec%, CL_R_Min, FI, OI, DF and ShEn.

### Rec% and CL_R_ are not correlated with myofiber anisotropy

Another tissue characteristic that has been shown to affect electrophysiological characteristics in atrial and ventricular tissue is myofibril orientation, with non-uniform myofiber orientation (i.e. greater anisotropy of fiber orientation) thought to be related to slow and inhomogeneous conduction.^3, 33, 34^ We therefore assessed the uniformity of fiber orientation in all 6 regions of the atria and assessed the relationship between myofiber anisotropy index (AI) and all electrogram measures. Figure 5A demonstrates varying degrees of fiber orientation in terms of AI in exemplary LAFW and LAA tissue sections. AI in the different regions in the atria is shown in Figure 5B. The highest AI (i.e. most uniform fiber orientation) was seen in RAFW and the lowest AI was seen in PLA. This regional variability of AI was different from the regional heterogeneity of Rec% and CL_R_ discussed earlier. Indeed, AI was not found to be correlated with Rec%, CL_R_ or other established EGM measures (FI, OI, DF and Shen; Figure 5C).

**Figure 5.**
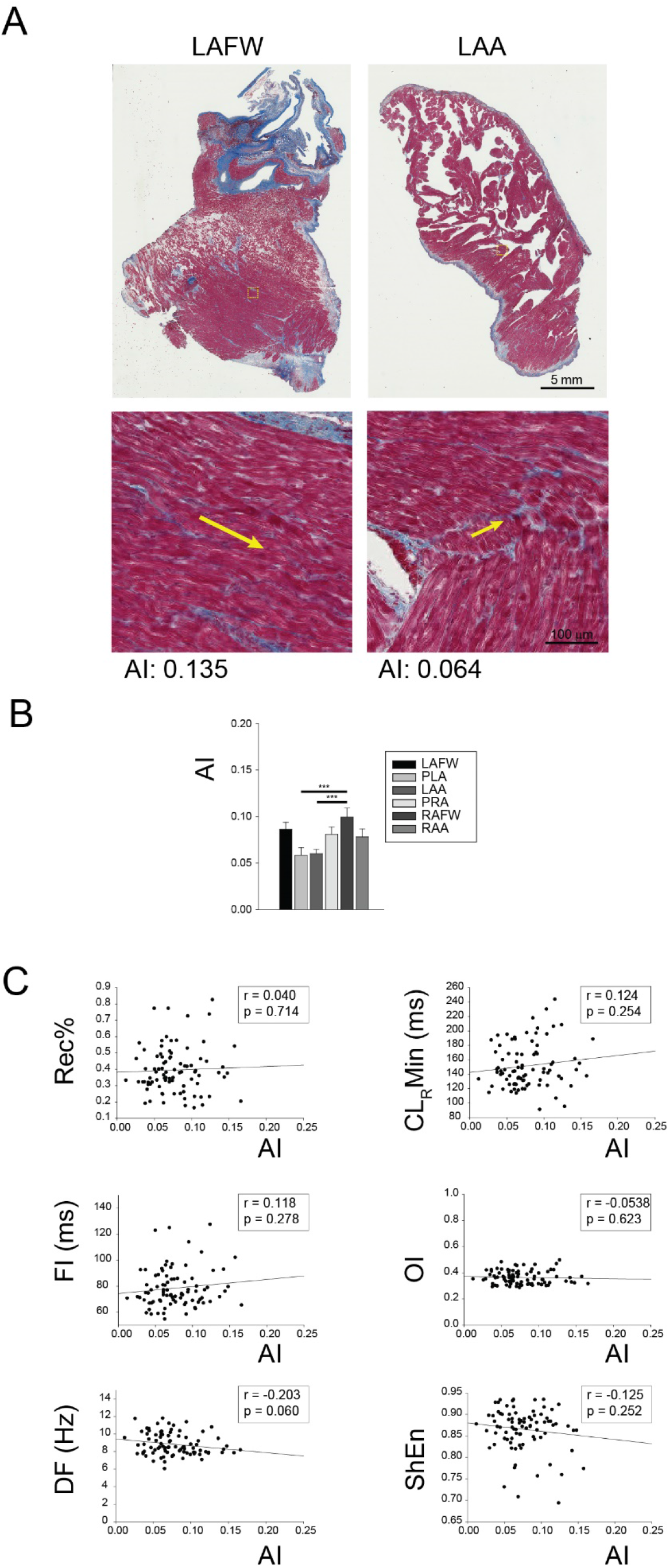
A, Example of fiber orientation measurements in LAFW and LAA. B, Regional differences in AI. Data are presented as mean ± SEM; *** p < 0.001. one-way ANOVA with Holm-Sidak method for pairwise multiple comparison. C, Correlation of AI with Rec%, CL_R_Min, FI, OI, DF and ShEn.

### Rec% is closely related to the spatial distribution of parasympathetic nerve fibers in the atria

Previous studies have shown significant parasympathetic nervous system remodeling in the atria, with the parasympathetic nervous system thought to contribute to the formation of substrate for reentry in the atria.13 We therefore examined the spatial relationship between parasympathetic innervation and EMR. We also examined the effects of parasympathetic blockade on EMR in each region of the left and right atrium.

We have previously shown that RAP leads to marked hypertrophy of parent nerve bundles in the PLA, resulting in a global increase in parasympathetic and sympathetic innervation throughout the left atrium^29^. Parasympathetic fibers were found to be more heterogeneously distributed in the PLA and LAFW when compared with the LAA^29^. In the current study, we assessed the relationship between spatial distribution of parasympathetic nerve fibers and underlying EMR. Figure 6A shows an example of significant parasympathetic fiber heterogeneity in the left atrium (i.e. high standard deviation); in contrast, figure 6B shows another region in the left atrium with significantly more homogeneous parasympathetic fiber distribution (i.e. lower standard deviation). We discovered that heterogeneity of distribution of parasympathetic nerve fibers was closely correlated with absolute value and the spatial heterogeneity of Rec% in the RAP atria (figures 6C and D respectively). Such a correlation with parasympathetic nerve distribution was not noted with any other AF electrogram parameter (Supplemental figure X). Next, we examined the effect of parasympathetic blockade on AF electrograms. With atropine, there was a significant decrease in Rec% in the LAFW (figure 6E). Changes in CL_R_ were more pronounced, with atropine leading to a significant increase in CL_R_ in the LAFW, but a significant decrease in CL_R_ in the RAFW and RAA (figure 6F). While the reasons for the differences in direction of change of CL_R_ between the left and right atrium are not clear (also see Discussion), these data when taken together indicate that parasympathetic innervation strongly influences recurrence morphology in the fibrillating atrium, with parasympathetic signaling leading to a significant change in CL_R_ both atria.

**Figure 6.**
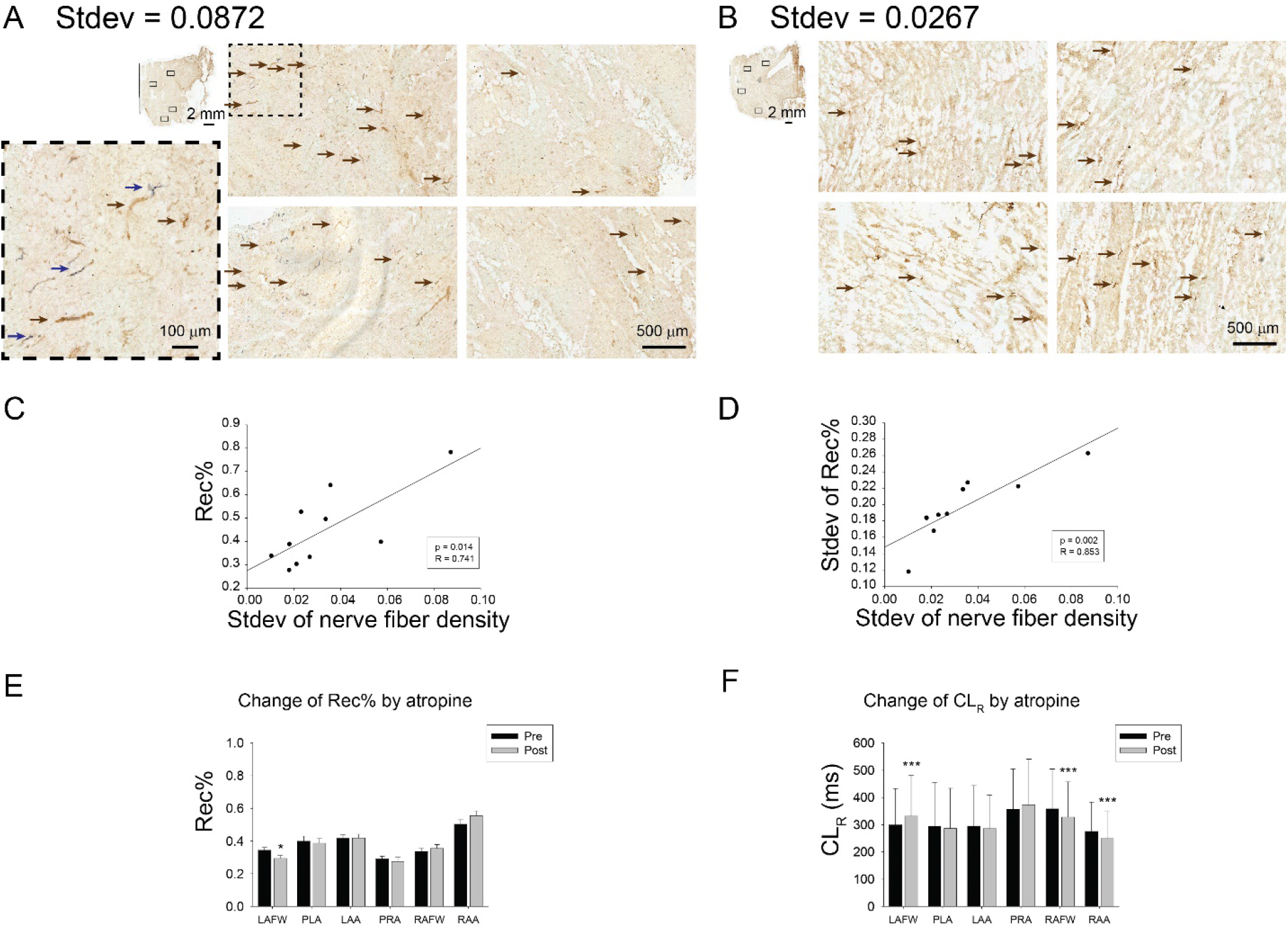
A and B, Representative micrographs of atrial regions with high (A) and low (B) standard deviation of parasympathetic nerve fiber density. Location of four random micrographs are denoted in mini map. Blue and brown arrows designate sympathetic and parasympathetic nerve fibers, respectably. C and D, Correlation of Rec% (C) and standard deviation of Rec% (D) with standard deviation of parasympathetic nerve fiber density. E, change of Rec% by atropine in six atrial regions. Data are presented in Mean ± SEM; * p < 0.05. paired t-test. F, change of CL_R_ by atropine in six atrial regions. Data are presented in Mean ± standard deviation; *** p < 0.001. paired t-test.

## Discussion

In our high-density biatrial epicardial mapping study in a canine model of persistent atrial fibrillation, we evaluated electrogram recordings in each of six atrial sub-regions and performed a comprehensive analysis of bipolar atrial electrograms.

Our results demonstrated that (1) the recurrence morphology measures Rec% and CL_R_ showed significant inter-regional differences across the different atrial sub-regions, with Rec% being greatest in the appendages and CL_R_ being lowest in the PLA; (2) across animals and regions, Rec% and CL_R_ correlated moderately with measures of AF fractionation (FI) and complexity (ShEn) but not with DF; (3) Rec% closely reflects stability of rotational activities (arrhythmogenic substrate) in the fibrillating atrium; (4) Rec% and CL_R_ had no significant correlation with atrial fibrosis and myofibril orientation; (5) the spatial distribution of Rec% corresponded closely to the spatial distribution of parasympathetic nerves, with CL_R_ demonstrating significant responsiveness to parasympathetic blockade.

### Limitations of previous attempts to find ‘order’ amongst complex AF activation patterns using AF electrograms

The detection of arrhythmogenic regions during AF is challenging. The dynamics of AF are complex and not clearly understood. Electrogram morphology is dependent on complex activation wavefronts during atrial fibrillation. Moe et al hypothesized that multiple, simultaneous reentrant depolarization wavefronts circulate in the atria.^35^ However, previous studies from Cox et al. and also from Konings et al during intraoperative studies of AF showed that the process of AF activation in isochronal maps is not random.^36, 37^ Further studies from Wells et al., Ropella et al. and Gerstenfeld et al. confirmed that wave-front propagation during AF is nonrandom by analyzing similarities between electrogram signals and by using the coherence spectrum.^38–40^ In a related study, Botteron et al. analyzed the spatial organization of AF and found that the correlation in sequences of activation decreased with the distance between the recordings exponentially with a higher correlation in paroxysmal compared to chronic AF.^41^ As a result, several investigators have postulated that detailed examination of the frequency, complexity and more recently morphology characteristics of AF electrograms may help determine the presence of potential AF driver sources, as well as yield information on the nature of electrical and structural remodeling in AF.^42, 43^ However, clinical attempts at using AF electrograms to detect and eliminate arrhythmogenic source regions (by using ablation) have had mixed results.^44^

Nademanee et al demonstrated a high AF success rate after ablation at regions that demonstrated CFAE.^44^ However, similar success rates have not been reproducible, with a recent randomized trial (STAR AF 2) demonstrating no benefit of additional CFAE ablation compared to pulmonary vein isolation alone in patients with persistent AF^45^. One reason for this seeming failure of CFAE ablation is that the pathological basis of CFAE in AF is still not clear. Although numerous studies have demonstrated that progressive atrial remodeling with AF persistence is associated with increasing atrial substrate complexity including electrogram fractionation, these studies used CFAE bipolar algorithms which were highly variable. Furthermore, most of these bipolar algorithms were found to correlate poorly with markers of AF substrate complexity such as conduction velocity, number of waves or breakthroughs per AF cycle, and electrical dissociation.^46^

Another electrogram measure that has evoked significant interest is cycle length. Regional differences in cycle length were reported in previous studies.^47^ These findings increased the interest in the analysis of the frequency spectrum as a measure of the activation rate in animal models.^43, 48–50^ Measurements of AF cycle length and frequency domain were used to guide ablation in patients with AF using high-resolution analysis of the Fourier power spectrum with its dominant frequency (DF).^51–56^ However, no beneficial effect of DF ablation compared to PVI alone has been convincingly demonstrated to date.^56, 57^ One measure of AF frequency that has shown some clinical promise is Organization Index (OI). It has been shown that AF episodes with high OI are more easily terminated with burst pacing and defibrillation.^20^ Jarman et al showed that at sites of organized activation, the activation frequency was also significantly more stable over time.^58^ This observation is consistent with the existence of focal sources, and inconsistent with a purely random activation pattern. Ablation of such regions was associated with organization of AF in remote atrial regions in patients using left atrial noncontact mapping. However, this study approach was limited to focal sources.

Ganesan et al showed that ShEn – a marker of signal amplitude distribution - may be associated with the pivoting zone of rotors in some cases.^59^ They showed that ShEn could differentiate the pivot from surrounding peripheral regions and thereby assist in clinical rotor mapping. However this method was limited to the pivot point of reentry, which might be detected only within 2-3 mm distance to rotor core.

Most recently, ablation strategies have targeted AF rotors as detected by the Focal Impulse and Rotor Modulation (FIRM) method.^60^ Using a basket catheter with 64 electrodes, this method provided a panoramic activation map and initially reported improved ablation outcome compared with conventional ablation alone.^60^ However, subsequent studies have not shown significant success with FIRM mapping in patients with persistent AF.^61^ Furthermore, it has been felt that the phase map algorithms may lead to possible over-detection of AF sources^62^.

Taken together, nearly all the electrogram based ablation strategies in AF – several of which have used FFT-based analyses of the AF electrogram - have had significant shortcomings. Below we discuss why novel electrogram morphology measures – Rec% and CL_R_ - may be superior to more established electrogram measures of AF at determining arrhythmogenic substrate for AF.

### Measures of EMR - Rec% and CL_R_ - may be superior to traditional AF electrogram measures in detecting AF sources

In AF electrograms the relative timings and morphologies are constantly changing. Indeed, recent studies have argued that the shape (morphology) and the repeatability of the electrogram signal over time provides significant information that is not contained in more typical frequency and complexity measures of AF. A recent study showed that a novel frequency analysis algorithm and longer duration of AF electrograms in search for temporally stable AF drivers have some promise^63^. Ciaccio et al showed that in paroxysmal AF, CFAE repetitiveness is low and randomness is high outside the PVs, particularly the left superior PV, and that in persistent longstanding AF CFAE repetitiveness becomes more uniformly distributed at disparate sites, possibly signifying an increasing number of drivers remote from PVs.^64^ Ciaccio et al further showed that the dominant repetitive electrogram morphology of fractionated atrial electrograms has greater temporal stability in persistent as compared with paroxysmal atrial fibrillation^65^. Our group recently developed a new electrogram measure which analyzes EMR,^16^ using modification of a method by Eckmann et al.^17^ Rec% describes the percentage of the most common morphology, with CL_R_ signifying the mean cycle length of activations of the most recurrent morphology.^66^

In this study, we discovered that discrete morphology patterns exist in AF and can be identified with the novel morphology recurrence plots. Rec% and CL_R_ are only somewhat correlated with established electrogram measures of AF fractionation (FI) and complexity (ShEn), and provide new information in quantifying the degree of repeatability of electrogram morphologies. We therefore believe that these new measures provide more information about the nature of arrhythmogenic AF substrate than more traditional electrogram measures for detecting AF sources. To test this hypothesis, we performed a systematic analysis of the number, stability and cycle length of 360 degrees rotational activity in different regions of the atria. Previous studies have found a strong relationship between the presence and number of these rotational activities and the ability of the atria to sustain AF^67^. In the current study, the stability of rotational activities in different sub-regions of the atria was found to correlate closely with Rec%. These data serve as an important initial validation of our postulate that sites of high recurrence morphology with the shortest cycle lengths – i.e. regions of low CL_R_ – may represent sites of AF drivers. These data also support further testing of this hypothesis in patients with persistent AF, by performing targeted ablation at sites of low CL_R_.

### Pathophysiological basis of sites of high morphology recurrence – role of parasympathetic nerve distribution in genesis of sites of high Rec% and low CL_R_

Since myofiber orientation and fibrosis are thought to be important contributors to the creation of arrhythmogenic substrate for AF,^34^ we systematically assessed the relationship between recurrence morphology measures and underlying myofiber orientation/fibrosis. We found that the appendages exhibited the highest Rec% compared to other parts of the left and right atrium. Previous investigations have suggested that myocyte fiber orientation may affect AF organization.^68^ However, we discovered no clear relationship between Rec% and myocyte fiber orientation in this study, indicating that other mechanisms may underlie this regional predilection of high Rec% for the appendages. In contrast to Rec%, CL_R_ was lowest in the PLA in the majority of animals. This is consistent with our initial clinical findings in patients with persistent AF^9^, where CL_R_ was the lowest in the PVs or PLA in nearly two thirds of all patients with AF (see below).

Some previous studies have attempted to relate EGM parameters such as voltage, fractionation, and DF with tissue characteristics like fibrosis. Marrouche et al. showed that there is a correlation between atrial fibrosis – as determined by delayed enhancement on MRI - and low-voltage regions.^69^ A related study suggested that CFAEs also correlate with regions of atrial fibrosis.^70^ However, a limitation of these studies was that they were not performed with high-resolution contact mapping; furthermore, detailed microscopic tissue analyses of fibrosis and anisotropy were not performed. In this study, we systematically analyzed several electrogram measures simultaneously with high-resolution mapping during AF, and then performed detailed tissue correlations with fibrosis. While the amount of fibrosis was discovered to be the least in the appendages, we discovered only a weak correlation between AF electrogram measures – both established measures as well as new morphology measures - and fibrosis.

An important upstream mechanism that is thought to contribute to electrical remodeling is increased activity of the parasympathetic nerve.^13^ We and others have shown in recent years that increased parasympathetic nerve sprouting – and a resulting increase in parasympathetic signaling in the atrium – is an important mechanism that contributes to electrical remodeling in the atrium.^15^ In a recent publication^29^, we showed that rapid atrial pacing leads to marked hypertrophy of parent autonomic nerve bundles in the PLA, resulting in a global increase in parasympathetic and sympathetic innervation throughout the LA. Parasympathetic fibers were found to more heterogeneously distributed in the PLA and LAFW when compared with the LAA. The coefficient of variation of CL_R_ was also found to be significantly greater in the PLA and LAFW than in the LAA indicating, suggesting that the spatial distribution of parasympathetic nerve fibers likely impacts recurrence morphology. In the current study, we assessed the precise relationship between spatial distribution of parasympathetic nerves and EMR. We discovered that the spatial distribution of parasympathetic nerve fibers was more closely related to Rec% than to any other electrogram parameter. Furthermore, parasympathetic blockade led to a significant change in CL_R_ in sub-regions of the right and left atrium, again demonstrating the significant contribution of parasympathetic signaling to EMR. Interestingly, the direction of change of CL_R_ in response to parasympathetic blockade differed between the right and left atrium. The mechanisms underlying these regional changes in CL_R_ may reflect differences in the precise pattern of parasympathetic innervation, M_2_ receptor and IK_Ach_ concentrations between the atria^71, 72^ and need to be further investigated in future studies.

### Study Limitations

Our animal model demonstrated smaller amount of fibrosis, mainly < 20%, than in previous studies that have attempted to correlated AF EGMs with underlying atrial fibrosis. It is possible that a stronger correlation between Rec% and fibrosis exists, in the presence of greater degrees of fibrosis. Further investigations therefore need to be performed with models incorporating a higher degree of fibrosis (30-40%). We detected fibrosis percentage only near the epicardial surface. Further studies showing differences of endocardial and epicardial fibrosis and EGM measures need to be conducted. The analysis of the parameter Rec% was calculated based on ten-second electrogram recordings in duration and may miss longer-term recurrence patterns. Further investigations of the dependency of Rec% on electrode size and geometry as well as distance to the tissue, near and far-field effects and endocardial vs. epicardial mapping need to be explored. Further investigations of the temporal stability of these new measures in paroxysmal and persistent AF need to be investigated.

## Conclusion

Traditional anatomically-guided ablation and attempts in the last decade to perform electrogram guided AF ablation (CFAE, DF, FIRM) have not been shown to be a sufficient treatment for persistent AF. We have extensively studied the mechanistic basis of a new electrogram guided approach to AF that combines high morphology recurrence with fast cycle length. Our results suggest that EMR parameters – Rec% and CL_R_ – may be more reflective of arrhythmogenic substrate for AF than any previously studied electrogram parameter of AF. Further studies are necessary to determine the effectiveness of this novel electrogram approach in guiding catheter ablation of persistent AF.

## ABBREVIATION

AF: Atrial fibrillation
AI: Anisotropy index
ANOVA: Analysis of variance
CFAE: Complex fractionated atrial electrograms
CL_R_: Cycle length of the most recurrent electrogram morphology
DF: Dominant frequency
EGM: Electrogram
EMR: Electrogram morphology recurrence
FI: Fractionation interval
LAT: Local activation time
LAA: Left atrial appendage
LAFW: Left atrial free wall
PLA: Posterior left atrium
PRA: Posterior right atrium
RAA: Right atrial appendage
RAFW: Right atrial free wall
Rec%: Recurrence percentage
PV: Pulmonary vein
SD: Standard deviation
SEM: Standard error of the mean
ShEn: Shannon’s entropy

